# Harnessing CRISPR/Cas12a Technology to Combat Wheat Dwarf Virus Without Genetic Modification

**DOI:** 10.1101/2024.07.01.598556

**Authors:** Buse Baran, Tarık Teymur, Ali Ferhan Morca, Muhsin Konuk, Cihan Taştan

## Abstract

Wheat (*Triticum aestivum* L.) is a staple crop globally, but its yield and quality are significantly affected by viral diseases, particularly the Wheat Dwarf Virus (WDV). Developing antiviral strategies is crucial for enhancing wheat productivity. This study aims to investigate the use of CRISPR/Cas12a as an antiviral agent against WDV without the need for genetic modification of the host plants. WDV genomic DNAs were isolated from wheat and barley samples collected from various regions in Turkey and other countries. Guide RNAs (gRNAs) targeting conserved regions in the coat protein (CP) gene of WDV were designed and synthesized. The CRISPR/Cas12a ribonucleoprotein (RNP) complex was tested for its ability to induce DNA cleavage in WDV genomes through in vitro assays, and results were analysed using agarose gel electrophoresis. The designed gRNAs effectively induced DNA cleavage in WDV genomes, with gRNA1 showing consistent performance across all tested isolates. gRNA2 exhibited variability, successfully targeting seven out of ten isolates. No off-target effects were observed in control assays, confirming the specificity of the gRNAs. The CRISPR/Cas12a system, guided by specifically designed gRNAs, can effectively target and cleave WDV genomes, providing a potent, non-GMO antiviral strategy. This approach has potential applications for managing WDV infections in wheat and barley, highlighting its versatility and effectiveness.

## INTRODUCTION

Grains, which are the raw material of daily bread in human nutrition, are widely used in animal nutrition and industry. Wheat (*Triticum aestivum L*.) is one of the most cultivated cereals in the world. Grain of wheat, which has a wide adaptation ability, is the staple food of approximately 50 countries due to its appropriate nutritional value, ease of storage and processing (Kün 1988). According to the data of the United Nations Food and Agriculture Organization (FAO), world wheat production is approximately 784 million tons in 2023. Wheat production is threatened by diseases caused by different factors every year, and as a result, yield and quality decrease significantly. Increasing air temperatures with the effect of global warming in recent years cause epidemics of some viral diseases carried by aphids.

Among the viral diseases detected by studies on grains, yellow dwarf virus diseases: Barley yellow dwarf viruses (BYDV-PAV, BYDV-MAV, BYDV-RMV, BYDV-SGV), Cereal yellow dwarf virus-RPV (CYDV-RPV) and wheat dwarf virus (WDV) is one of the important viral diseases that cause yield and quality losses (Pocsai et al. 2003, İlbağı 2003, İlbağı 2006, İlbağı et al. 2006). WDV is a species belonging to the genus Mastrevirus. The mastrevirus has a monopartite single-stranded circular DNA genome, and the typical mastrevirus genome encodes four different proteins: movement protein (MP), coat protein (CP), and two replication-associated proteins (Rep and RepA). The presence of an intron in the rep gene enables mastreviruses to produce two different forms of the replication protein. The long intergenic region (LIR) and short intergenic region (SIR) contain sequence elements required for viral replication and transcription (Kis et al 2016).

Antiviral agents need to be developed to increase wheat productivity. Thanks to the developing technology, the CRISPR (Clustered regularly interspaced short palindromic repeats)/Cas system has been adapted to precisely edit the genomes of many species (Nekrasov et al. 2013). CRISPR associated (CRISPR-Cas) is an adaptive immune system in prokaryotes that protects against invading bacteriophages by cleaving the DNA of bacteriophages (Horvath and Barrangou 2010; Garneau et al. 2010; Gasiunas et al. 2012). Successful examples of regulation in different plant species include rice, maize, wheat, soybeans and tomatoes (Mikami et al. 2016; Lee et al. 2019; Kelliher et al. 2019; Biswas et al. 2019; Svitashev et al. 2016); Gil-Humanes et al. 2017; Okada et al. 2019; cai et al. 2019). Three main types of CRISPR systems have been described so far: Types I, II, and III. Of the type II CRISPR systems, CRISPR-Cas9 and CRISPR-Cas12a are the two main nucleases used to edit plant genomes (Nekrasov et al. 2013; Svitashev et al. 2015; Kim et al. 2017). The CRISPR-Cas9 system recognizes an NGG sequence called PAM (protospacer adjacent motif) and creates double helix breaks (Svitashev et al. 2015; Kim et al. 2017). On the other hand, the CRISPR-Cas12a system recognizes the TTTV PAM sequence and creates double-strand breaks in a different region, ie downstream. PAM recognition sites offer different options for gene editing in different parts of the genome. Cas9 is an efficient option for regulating GC-rich regions, while Cas12a performs better in AT-rich regions. Also, the intersecting results of Cas9 and Cas12a differ. Cas9 creates sharp-edged DNA breaks close to the PAM region, while Cas12a creates gradual breaks further away from the PAM region (Svitashev et al., 2015; Kim et al., 2017). These different cut characteristics offer different uses in gene editing applications.

All studies in the literature have been achieved by modifying the plant genome or by performing mutagenesis studies. In this study, we aimed a CRISPR system that can bind directly to the virus without genetically modifying the plants, and we performed a proof-of-concept study showing whether it can target the WDV genome as an antiviral strategy using virus-specific CRISPR/Cas RNP structures without integrating into the plant genome. In this study, we have shown that the designed sgRNA sequences can effectively produce DNA breakage on the WDV genomes isolated from the virus from different cities in Turkey. This data suggests that Cas12 plus sgRNA ribonucleic protein complect can effectively target WDV genomes. And this study proves us to use CRISPR/Cas RNP complex in the agricultural field to test WDV infection in wheat.

## MATERIALS AND METHODS

### Sample Collection and WDV genome extraction

Wheat Dwarf Virus genomic DNAs were isolated from wheat and barley collected from different countries (Hungary_JQ647455-AM040732, Sweden_ NC_003326, Iran_FJ620684-JN791096) in the NCBI database and from different cities (OQ183228_KRSH-I43, OQ183229_KRSH-I66, OQ230451_KRSH-I41, OQ190468_KRSH-I38, OQ183230_KNY-I76, MW387505_KNY-I34, MW387502_AFY-I54, MW381791_NVS-I56) in Turkey, using the previously mentioned method (Morca et al. 2021). Samples were collected randomly in the vicinity of winter barley and wheat fields with suspected WDV infection from 10 different locations in 6 six provinces (Afyon, Ankara, Kırşehir, Konya, Nevşehir, and Yozgat) (Morca et al. 2021). Each location was considered as one sample and 10 samples (6 barley and 4 wheat) were collected. The total DNAs of 10 samples were extracted from wheat and barley plants using the DNeasy Plant Mini Kit (Qiagen, Hilden, Germany, Cat No./ID: 69104) according to the manufacturer’s instructions.

### gRNA design and synthesis for Cas12a

Multiple sequence alignment of WDV DNA sequences was performed using the Clustal Omega. As a result of the alignment, conservative regions in the coat protein region of WDV were detected. Two different guide RNAs were designed based on these conserved regions. Guide RNA sequences “5-CGUGUCACGACGGAGUGGAU-3” and “5-AUGUUGUAUGUGCCUAUACG-3” were synthesized by GenScript by adding the Cas12a scaffold sequence (5-UAAUUUCUACUCUUGUAGAU-3) to the 5-base upstream region.

### Targeting analysis of CRISPR/Cas12a RNP against WDV genome isolates

Guide RNAs were dissolved by adding 314 μL of deionized water to a final concentration of 10 ng/μL. 50 μg of GenCRISPR Cas12a (Cpf1) Nuclease (Genscript, Cat. No: Z03502, USA) was dissolved in 1000 μL (100 μL of 10X Cas12a Nuclease Reaction Buffer and 900 μL of deinozied water). 10X Cas12a Nuclease Reaction Buffer was prepared to pH 7.9 with 500mM NaCl, 100mM Tris-HCL, 100mM MgCl2 and 1mg/mL BSA. A 20 μl reaction in 1xCas12a Nuclease Reaction Buffer containing 60 ng WDV ssDNA, 10 ng gRNA, and 100 ng GenCRISPR Cas12a (Cpf1) Nuclease for 30 mins at 37°C results in a digestion efficiency of the linearized plasmid (for each WDV ssDNA). Samples loaded on the 1% agarose gel prepared in 1X TAE buffer solution using a BIO-RAD gel electrophoresis system were run at 100 V- and 30-min. Visualization, documentation, and analysis were then performed with Gel Doc™ XR+ (Serial Number 721BR16450, Software Version 6.0.1.34).

### Statistics Analysis

Two-tailed Homoscedastic t-tests were performed using SPSS software. Outliers were not excluded in any of the statistical tests and each data point represents an in-dependent measurement. Bar plots report the mean and standard deviation or the standard deviation of the mean. The threshold of significance for all tests was set at * p <0.05. ns: non-significant.

## RESULTS

### WDV Specific Dataset Preparation

In this study, the diversity of mutations in WDV genomes isolated from wheat and barley were investigated for an effective guide RNA design. WDV genome reference sequences were obtained from the NCBI database, and the reference numbers of these sequences are JQ647455, AM040732, NC003326, FJ620684, JN791096, respectively **(Figure 1)**. In addition, the reference numbers of WDV genomes isolated from wheat and barley collected by us from Kırşehir, Konya, Afyon and Nevşehir in Turkey are OQ183228, OQ183229, OQ230451, OQ190468, OQ183230, MW387505, MW387502, MW381791, respectively (**Figure 1**).

**Figure 1:**
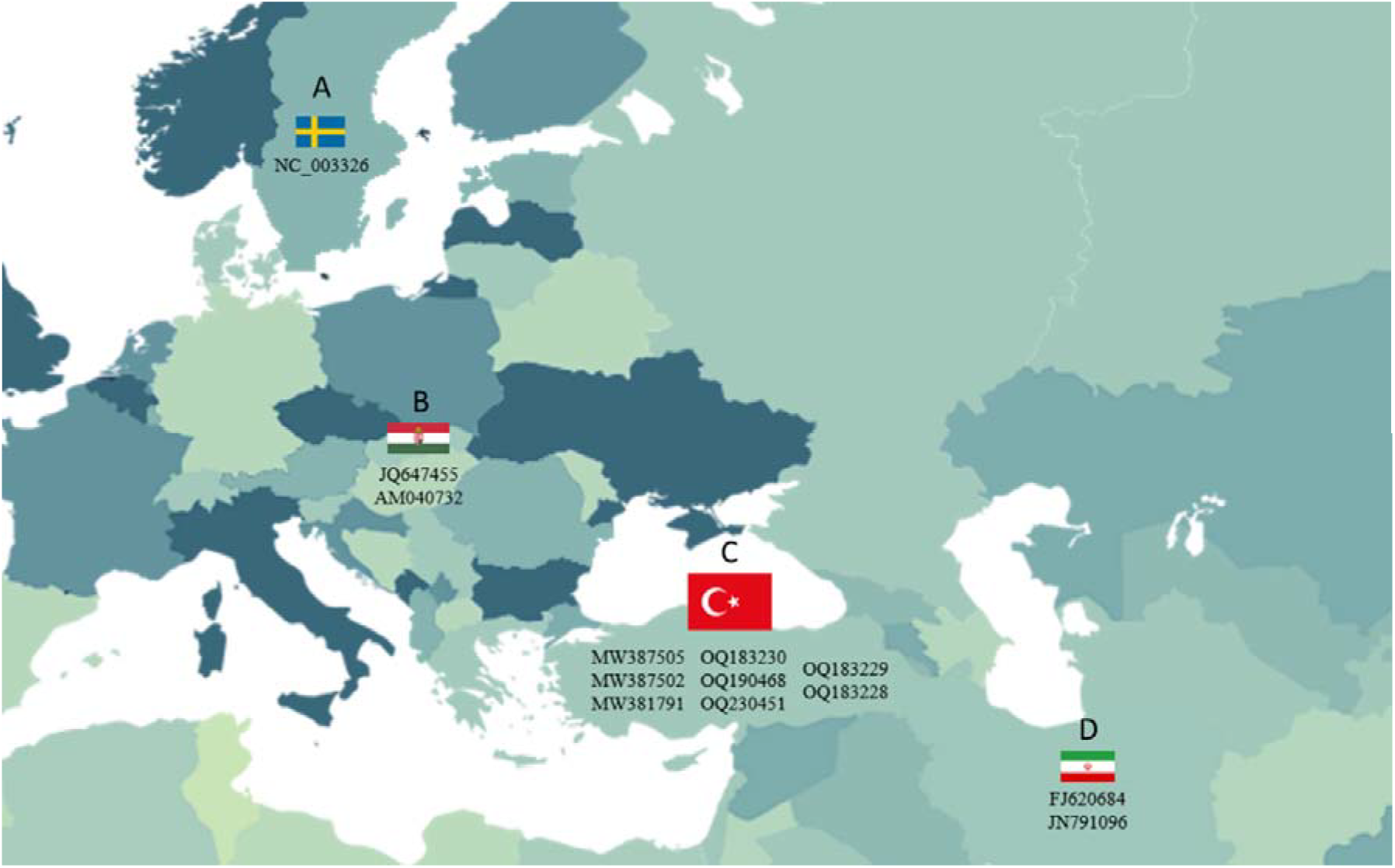
WDV strains isolated from different countries by NCBI sequence number (NC003326), GenBank numbers (JQ647455, AM040732, FJ620684, JN791096, MW387505, MW387502 and MW381791) and others (OQ183228, OQ183229, OQ230451, OQ190468, OQ183230). **A**.Sweden, **B**.Hungary, **C**.Turkey, **D**.Iran.

### Global Multiple Alignment of WDV and Target Selection

WDV is a virus persistently transmitted by dwarf cicadas in *Psammotettix alienus* and *P. provincialis* species in the Cicadellidae family and infects wheat and barley plants (Abt, I.,2019). The CP protein mediates vector (*Psammotettix alienus*) transmission and is vector specific. The genetic material of the virus is stored in this protein structure. It also allows the virion to attach to the host (wheat or barley) and penetrate the host cell membrane. (Abt, I.,2019) Therefore, conserved sequences in the coat protein (CP) gene of WDV strains isolated and sequenced from four different countries (Figure 1) were detected. In this study, the CP protein of WDV was targeted, and the reference sequences obtained from the database and the sequences of the strains isolated by us were performed using Clustal Omega software **(Figure 2)**. As a result of the alignment, conserved sequences in the coat protein (CP) gene were detected **(Figure 2A)**. Single point mutations were observed in the region of conserved sequences **(Figure 2A)**. These single point mutations were taken into account while designing the guide RNA, and two guide RNAs containing different numbers of single point mutations were designed **(Figure 2A-B)**. Guide RNA 1 was designed to contain two single point mutations in the overall sequence, without any mutations in the PAM sequence. Here, T and C with the highest information content were selected **(Figure 2B)**. Guide RNA 2 was designed to contain one point mutation in one PAM sequence and one in the overall sequence **(Figure 2B)**. There is a T-C mutation in the PAM sequence, and the “TTTG” PAM sequence suitable for Cas12a was preferred. In the general sequence, T with the highest information content was chosen **(Figure 2B)**. In subsequent experiments, these designed guide RNAs were used.

**Figure 2:**
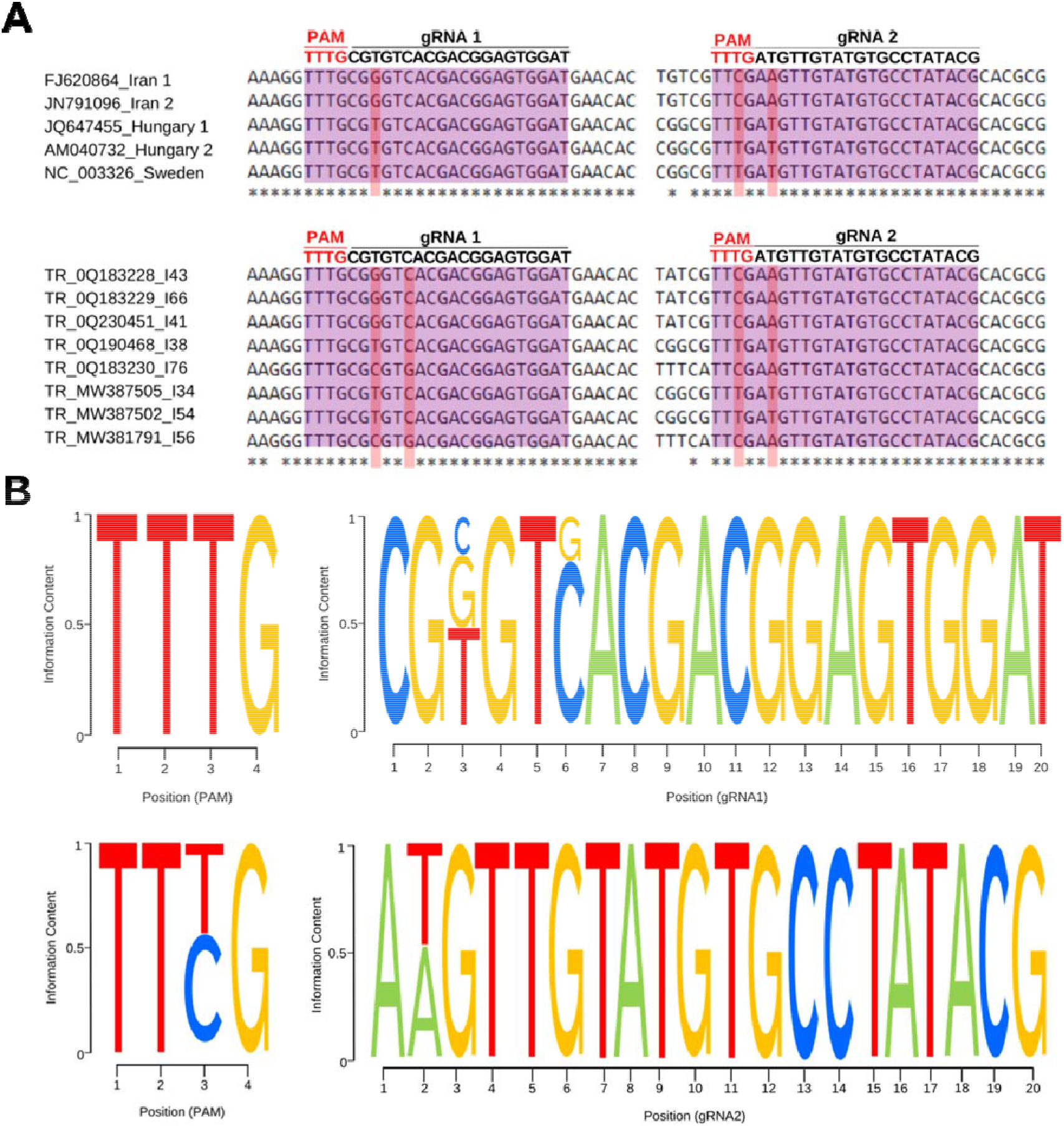
Multiple sequence alignment and information content of WDV genomes. **A**. Conserved regions in the CP protein gene and PAM sequences determined based on these regions, along with guide RNA designs, resulting from multiple sequence alignment. **B**. The frequency of variants in the targeted region of CP genes. Sequence logo representation of guide RNAs.

### Proof-of-concept for CRISPR/Cas System

Cas12a can be utilized as an antiviral agent because it can bind to single-stranded DNA and target the genomes of viruses. In this study, we aimed to test whether Cas12a, in conjunction with guide RNAs, can target isolated WDV genomes and induce DNA cleavage. For this purpose, we designed a proof-of-concept experiment. It is shown in principle in order to confirm that our theory has practical potential to demonstrate the viability of the work. First, guide RNAs designed specifically for WDV genomes isolated from wheat and barley were incubated with Cas12a, and then run on an agarose gel to evaluate the presence of CRISPR/Cas12a-related DNA breaks on WDV genomes **(Figure 3)**.

**Figure 3:**
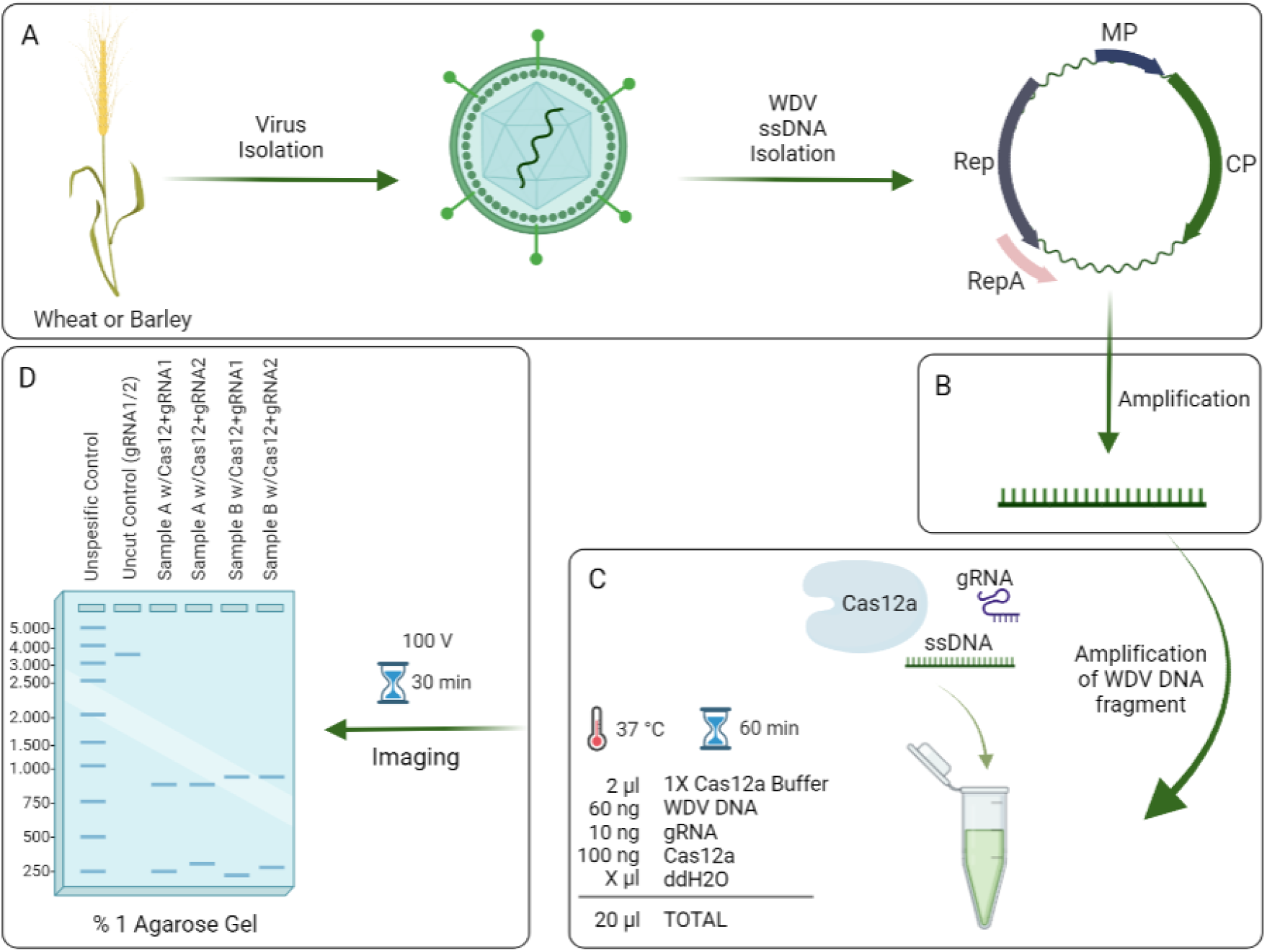
Experimental Set-Up of WDV specific Cas12a as an antiviral agent. **A**. WDV ssDNA genome isolated from wheat or barley. **B**. Amplification with CP primers using a thermal cycler. **C**. Co-incubation of WDV genomes with gRNA and Cas12a RNP complexes.**D**. Conducting gel electrophoresis after incubation.

### Study with CRISPR/Cas12a RNP structure on WDV

In order to detect DNA breaks that we can create in the targeted WDV genome with Cas12a, we aimed to determine the rates of DNA breaks with a special visualization method by running it on agarose gel. As a result of co-incubation, it was tested whether the designed guide RNAs cleaved in the WDV genome. We wanted to determine whether the two guide RNAs that form the RNP complex together with Cas12a will bind to unspecific circular DNA controls that do not contain the region we target, and whether they will create any unspecific breaks. In our first analysis for this, we showed that there is no such breakdown **(Figure 4A)**. Next, we wanted to test whether WDV genomes isolated from different cities could be broken down jointly. In 10 isolated WDV genomes, guide RNA 1 was observed to break through the whole WDV genome **(Figure 4B-4C and Supplementary file Figure 2**). It was observed that guide RNA 2 was able to break through seven WDV genomes, but not in three WDV genomes (MW387505, MW387502 and MW381791) **(Figure 4B and Supplementary file Figure 2)**. WDV genomes MW387505 and MW387502 match exactly with gRNA2, while WDV genome MW381791 has two different mutations (PAM: TTTG → TTCG and Seq: ATG → AAG) according to gRNA **(Figure 2A)**. These results showed us that two guide RNAs designed specifically for WDV could identify WDV genomes isolated from different regions in a statistically significant way and break DNA after binding. This suggests that Cas12a and WDV-specific guide RNAs can be used as antiviral agents **(Figure 4C)**.

**Figüre 4:**
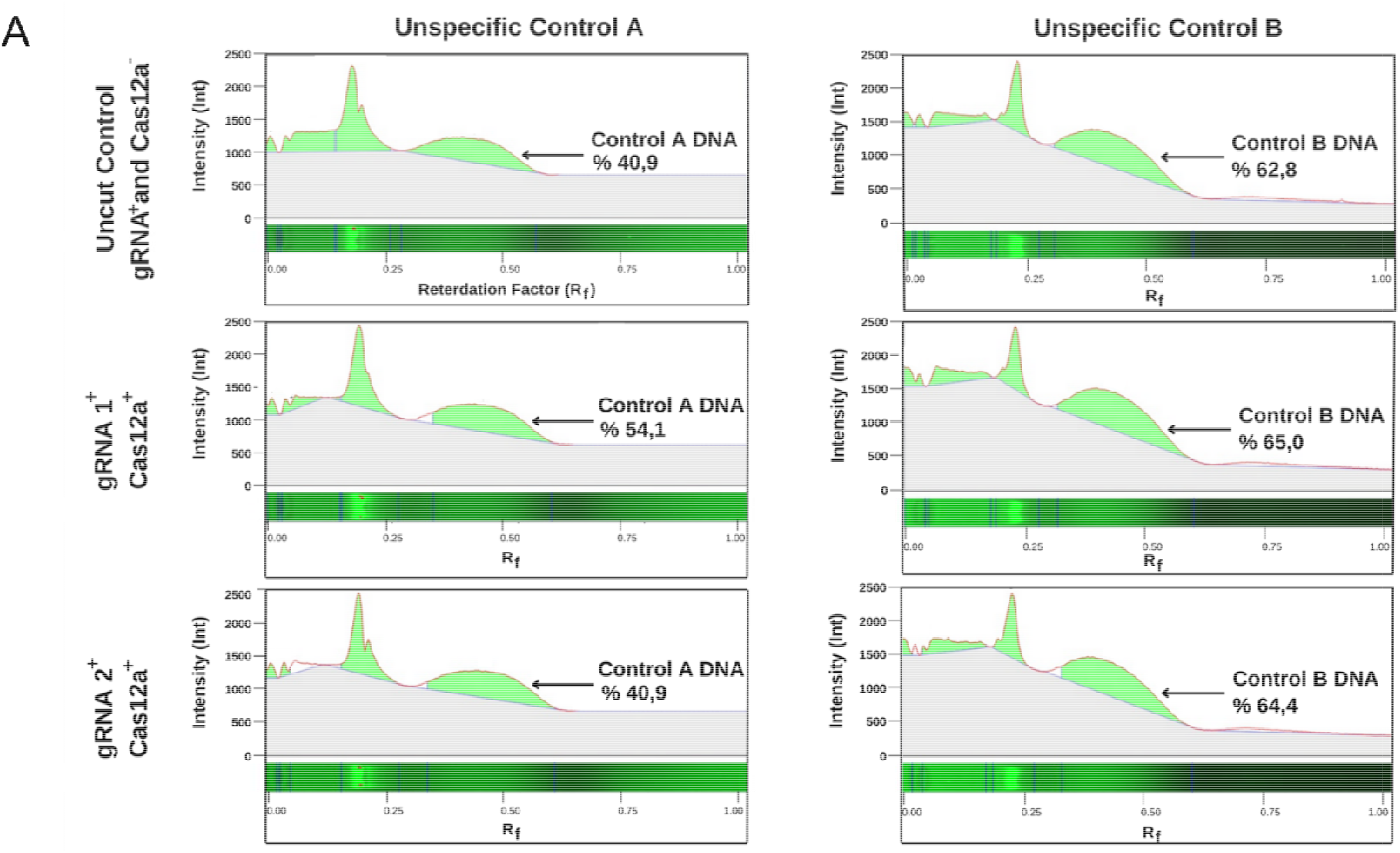
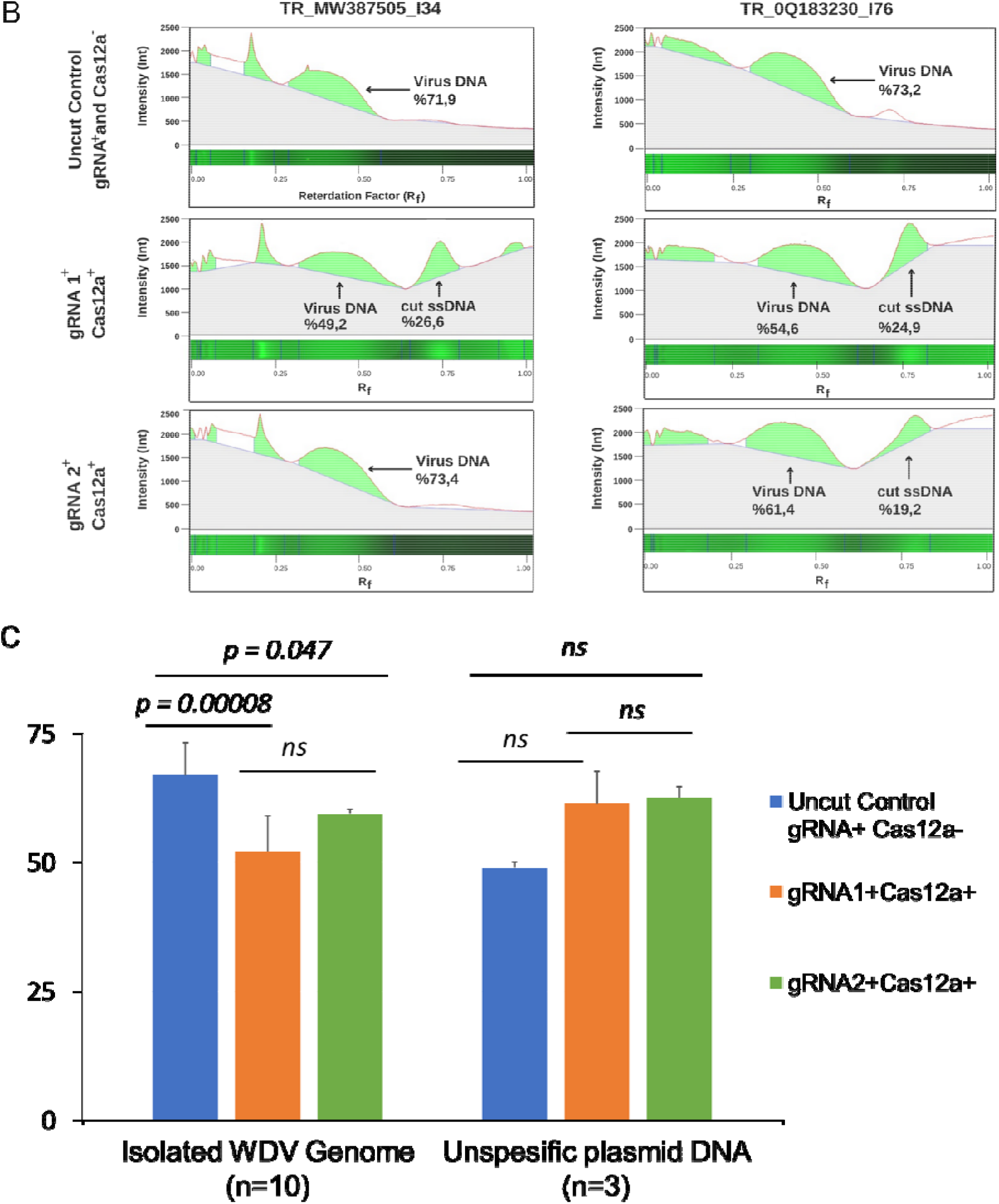
**A**. The unspecific control A and B plasmid DNAs were incubated in a Cas12a^-^, gRNA1^+^ & Cas12a^+^, and gRNA2^+^ & Cas12a^+^. Subsequently, after running on a 1% agarose gel, they were visualized using Gel Doc™ XR^+^. **B**. The TR_MW387505_I34 and TR_OQ183230_I76 Wheat Dwarf Virus (WDV) DNAs were incubated with uncut control (gRNA^+^ & Cas12a^-^), gRNA1^+^ & Cas12a^+^, and gRNA2^+^ & Cas12a^+^. Subsequently, after running on a 1% agarose gel, they were visualized using Gel Doc™ XR^+^. **C**. Statistical analysis of the comparison of results obtained from the incubation of WDV DNAs and unspecific DNAs with gRNA1 and gRNA2, as well as their incubation with uncut control.

## DISCUSSION

The application of the CRISPR/Cas12a system as an antiviral strategy against the Wheat Dwarf Virus (WDV) demonstrates a novel approach that does not require genetic modification of the host plants, thereby preserving their original genomic integrity. The effectiveness of CRISPR/Cas12a in targeting and cleaving WDV genomes was evident across various isolates from different geographic regions, indicating its potential broad-spectrum application (Kis et al., 2016).

In this study, we observed that the designed guide RNAs (gRNAs) showed a significant capability to induce DNA cleavage in WDV genomes, particularly with gRNA1, which was effective across all tested isolates. This finding aligns with previous studies that have highlighted the efficiency of CRISPR/Cas systems in editing viral genomes (Nekrasov et al., 2013; Lee et al, 2019). The ability of gRNA1 to function despite the presence of mutations suggests a robust targeting mechanism that can accommodate minor genetic variations within the WDV population, a critical feature for managing viral pathogens with high mutation rates (Kelliher et al., 2019). Conversely, gRNA2’s inability to induce cleavage in some WDV isolates indicates the importance of precise gRNA design, particularly in regions with potential single nucleotide polymorphisms (SNPs) that could affect binding efficiency (Kim et al., 2017). This specificity underscores the necessity of comprehensive genomic analyses to identify conserved viral regions that can serve as reliable targets for CRISPR/Cas12a interventions (Gil-Humanes et al., 2017).

The absence of unexpected results in the unspecific and uncut controls reinforces the specificity of the designed gRNAs, as no off-target effects were observed. This high specificity is a significant advantage of the CRISPR/Cas12a system, minimizing the risk of unintended genomic alterations in non-target organisms (Svitashev et al., 2016). Furthermore, the successful application of CRISPR/Cas12a as an antiviral agent without integrating into the plant genome presents a promising alternative to traditional plant genetic modification approaches. This method not only preserves the genetic integrity of the crop but also simplifies regulatory hurdles associated with genetically modified organisms (GMOs) (Nekrasov et al., 2013).

In summary, our findings indicate that the CRISPR/Cas12a system, guided by specifically designed gRNAs, can effectively target and cleave WDV genomes, providing a potent tool for managing WDV infections in wheat and barley. Future research should focus on optimizing gRNA design to enhance targeting efficiency and exploring the application of this system against other plant viruses to further validate its versatility and effectiveness (Cai et al., 2019; Okada et al., 2019).

## Acknowledgements

Figures of the study were drawn by B.B. and C.T. with BIORENDER.com.

