## Supplementary File for "Harnessing CRISPR/Cas12a Technology to Combat Wheat Dwarf Virus Without Genetic Modification"

**Table 1.** WDV isolates obtained from various regions in the NCBI database.

| **Isolate Name** | **Accession No** | **Refseq No** | **Host** |
| --- | --- | --- | --- |
| Iran/2008/B | FJ620684 |  | *Hordeum vulgare* |
| Iran/Bavanat/2010/D | JN791096 |  | *Hordeum vulgare* |
| Hungary/Kompolt10/  1/2010/C | JQ647455 |  | *Triticum vulgare* |
| Hungary/B/2005/E | AM040732 |  | *Triticum aestivum* |
| Sweden/wdv-[Enk1] |  | NC_003326 | *Triticum aestivum* |

**Table 2.** PCR Analysis Results of WDV Isolates (Morca F.A. ve diğerleri, 2021).

| **Isolate Name** | **Host** | **City** | **Yar** |
| --- | --- | --- | --- |
| KNY-I34 (MW387505) | Wheat | Konya | 2019 |
| KNY-I38-I39 (2 Samples) | Wheat | Konya | 2019 |
| KRSH-I41-I43 (2 Samples) | Barley | Kırşehir | 2019 |
| AFY-I54 (MW387502) | Wheat | Afyon | 2020 |
| NVS-I56 (MW38791) | Barley | Nevşehir | 2020 |
| ANK- I66-I76-I77 (3 Samples) | Barley | Ankara | 2020 |

**Figure 1.**

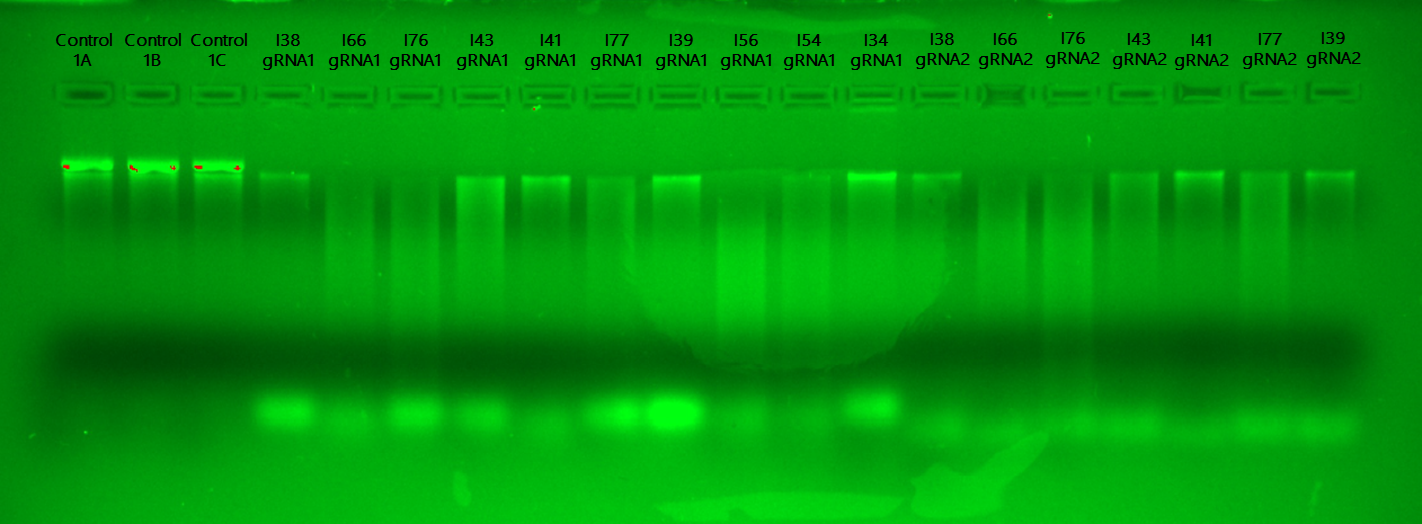

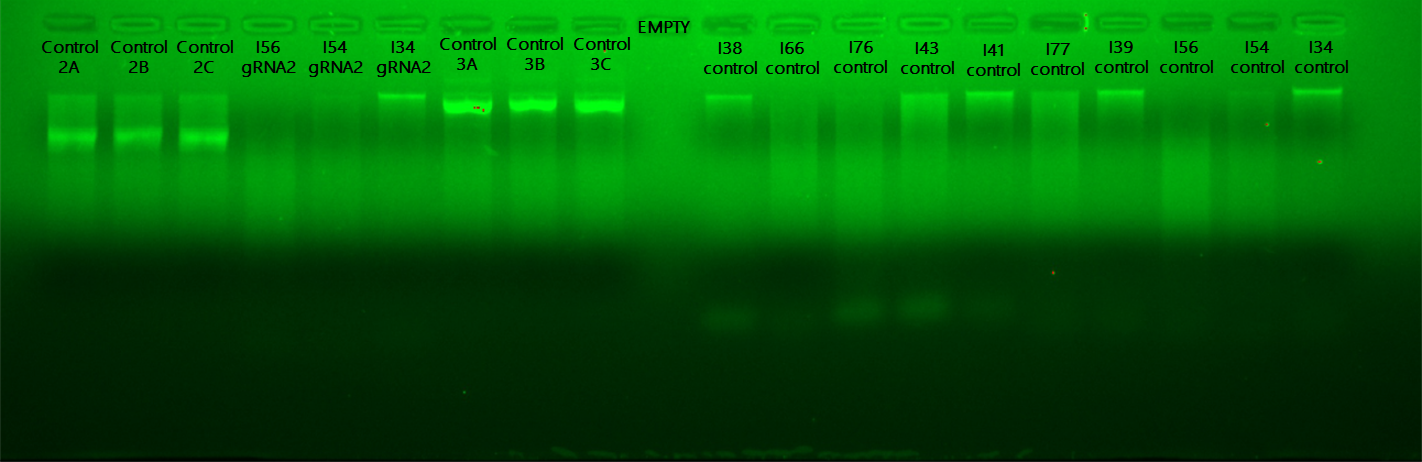

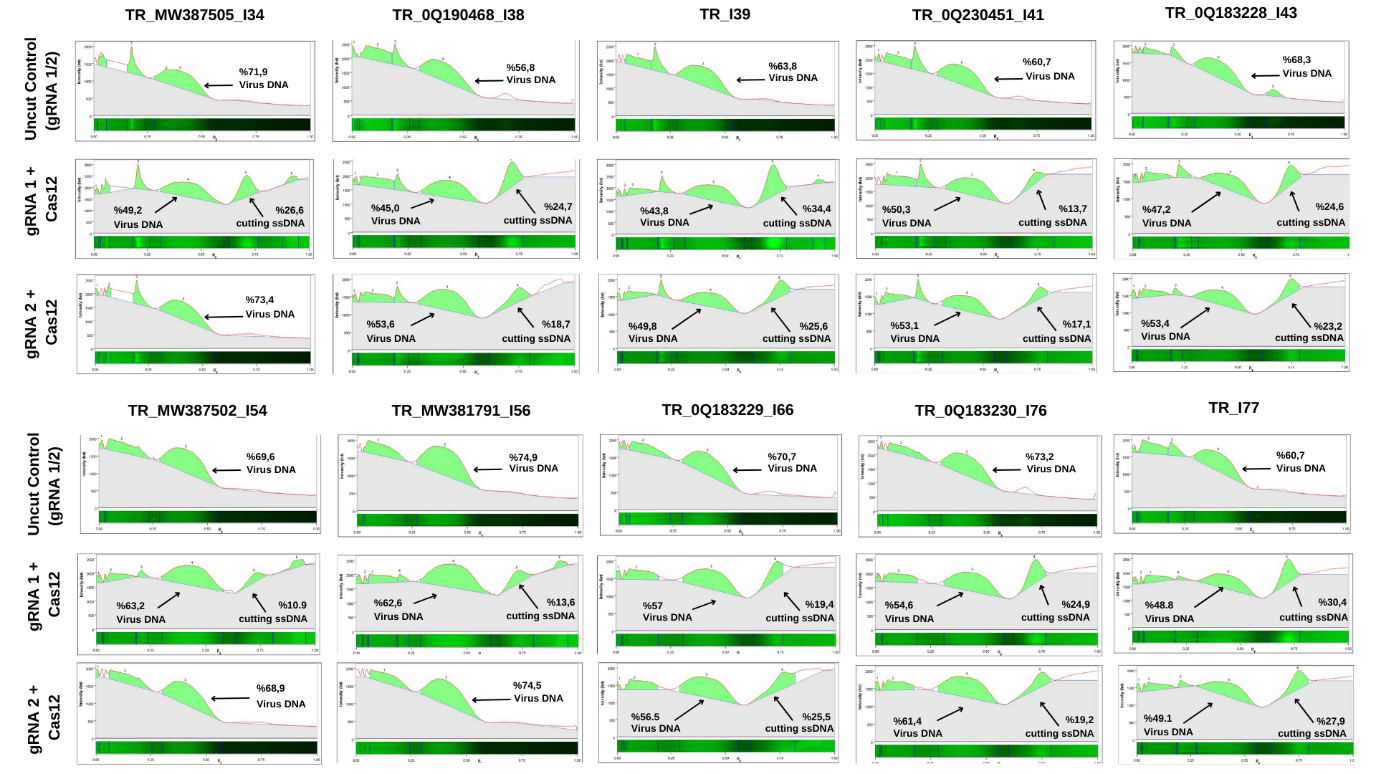

**Figure1.** The unspecific control A and B plasmid DNAs were incubated in a Cas12a^-^, gRNA1^+^ & Cas12a^+^, and gRNA2^+^ & Cas12a^+^. Subsequently, after running on a 1% agarose gel, they were visualized using Gel Doc™ XR^+^
